# Cataloging Viral Diversity from Nonaxenic Terrestrial Cyanobacteria Cultures

**DOI:** 10.1101/2022.12.14.520320

**Authors:** Cassandra L. Ettinger, Sudipto R. Paul, Neptali Flores, Ryan D. Ward, Nicole Pietrasiak, Jason E. Stajich

## Abstract

Viruses are exceedingly common, but little is known about their diversity let alone how they behave in extreme environments, and whether viruses facilitate adaptation of their hosts to harsh conditions. To set a foundation for understanding of these understudied viral-host interactions, we created a catalog of viruses through analysis of metagenomes from 50 unialgal but nonaxenic Cyanobacteria cultures with 47 cultures isolated from various terrestrial habitats, including desert soil and rock surfaces, tropical soil, and vernal pools. These cultures represent low diversity microbial consortia dominated by the terrestrial Cyanobacteria and its associated cyanosphere microbiome containing heterotrophic microbes. We identified viral sequences in metagenomes, grouped these into viral operational taxonomic units (vOTUs) and then placed vOTUs into viral clusters (VCs). We also calculated vOTU relative abundance and predicted possible bacterial hosts. In total we predicted 814 viral sequences representing 726 vOTUs. We assigned putative taxonomy to 72 of the 814 putative viral sequences; these were distributed into 15 VCs — mostly assigned to the recently abolished Caudovirales order (now Caudoviricetes class) of viruses. We found that nonaxenic cultures were dominated by unclassified and unclustered viral sequences. Furthermore, we predicted putative bacterial hosts for 211 vOTUs, with the majority of viruses predicted to infect a Proteobacteria (now Pseudomonadota) host. Overall, while limited, these results are consistent with the notion that both viruses and Cyanobacteria isolated from extreme environments are underrepresented in reference datasets. This work increases knowledge of viral diversity and sets a foundation for future exploration of viruses associated with terrestrial Cyanobacteria and their heterotroph associates, such as connecting specific viruses to critical cycling processes and investigating their metabolic functions.

## Introduction

Viruses infect all domains of life and are thought to be the most abundant biological entities on Earth (Paez-Espino et al., 2016). Viruses rely on their hosts for replication, and thus are intricately entwined with their host communities. In the lytic life cycle viruses infect their hosts, hijack the cellular machinery to replicate, and induce host cell lysis for dispersal of the newly created viral particles (Guttman, Raya & Kutter, 2004). As a host defense mechanism, phage infections can be evidenced in host’s genomes resulting from past infection events building up hosts’ phage immunity (Barrangou et al., 2007; Hampton, Watson & Fineran, 2020). Frequently viruses can also integrate and persist in their host’s genome as proviruses, with many bacteria harboring at least one prophage (Casjens, 2003). Prophages can have auxiliary metabolic genes (AMGs) or pathogenicity genes which they use to promote host growth (Bragg & Chisholm, 2008; Thompson et al., 2011; Fortier & Sekulovic, 2013; Hurwitz, Hallam & Sullivan, 2013). With such controls on host metabolism and fitness, viruses can have important and often underappreciated effects on bacterial population dynamics and turnover in microbial communities (Hurwitz et al., 2014; Hurwitz & U’Ren, 2016; Crummett et al., 2016).

Viral discovery has long been hindered by the lack of universal gene markers. However, modern machine-learning based and k-mer approaches have enabled researchers to robustly search metagenomes for viral sequences (Sullivan, 2015; Ren et al., 2017; Amgarten et al., 2018; Kieft, Zhou & Anantharaman, 2020; Guo et al., 2021). With these new methodologies, researchers can utilize available public sequencing data to discover and explore complex viral communities across previously uncharted ecosystems. However, virus discovery and potential host interactions and feedbacks to microbial communities in extreme environments, such as cave wall biofilms, biofilms of ephemeral waterfalls, temporal vernal pools, and dryland soils, are still notably understudied.

Cyanobacteria are keystone taxa of such polyextreme terrestrial environments due to their important roles in primary production and microbial organic matter build up, nitrogen fixation, bioweathering, and biofilm, microbial mat or biological soil crust formation (Paerl, Pinckney & Steppe, 2000; Perera et al., 2018; Pombubpa et al., 2020; Velichko et al., 2021). As essential ecosystem engineers, Cyanobacteria interact with a broad variety of other microbes. For examples, filamentous biological soil crust cyanobacteria house their own set of associated microbes established around the polysaccharide rich sheath material forming the cyanosphere, a microbial community similar to the copiotrophic rhizosphere microbiomes around plant roots (Couradeau et al., 2019). Cyanobacteria also influence free-living co-occurring microbes as shown in another study in which cyanobacteria in hypersaline microbial mats generated fermentation byproducts that promoted the metabolism of Chloroflexi and Sulfate-reducing bacteria (Lee et al., 2014). However, our knowledge on Cyanobacteria phages, viruses that infect Cyanobacteria, is still largely based on prior investigations of samples collected from aquatic ecosystems (Patzelt et al., 2014; Pietrasiak et al., 2019, 2021; Ward et al., 2021; Baldarelli et al., 2022). Cyanophages in aquatic ecosystems have shown to carry AMGs related to photosynthesis, phosphorus utilization, and carbon metabolism which are thought to aid phage fitness (Bailey et al., 2004; Sullivan et al., 2005, 2010; Millard et al., 2009). Yet, if Cyanophages have similar dynamic roles in polyextreme terrestrial habitats, or how phages might impact Cyanobacteria associated microbiomes is still virtually unknown.

Previously, Ward et al. (2021) generated 50 metagenomes from nonaxenic Cyanobacterial cultures, which represented unialgal polycultures containing primarily Cyanobacteria and heterotrophic microbial associates living in the cyanosphere. The majority of the polycultures (47/50) were obtained from Cyanobacteria isolated from terrestrial habitats, with over half representing cultures from arid environments (28/50) (Ward et al., 2021). Due to the paucity of information on phage and associated host identity for Cyanobacteria-dominated microbial communities of polyextreme environments, investigating low diversity communities such as cyanosphere microbiomes of cultured Cyanobacteria hosts is a tangible starting point of discovery. In this study, we surveyed the viral diversity associated with these Cyanobacterial co-cultures as part of a remote undergraduate research experience. We sought to build foundational knowledge about these viruses by (i) creating a catalog of viral genomes present in these co-cultures, (ii) assessing the relative abundance of viral genomes across co-cultures, and (iii) beginning to explore possible bacterial hosts for these viruses.

## Methods

### Data collection

Publicly available metagenomic samples, obtained from whole community DNA of Cyanobacteria-dominant nonanexic unialgal cultures and previously reported on in Ward et al (2021), were downloaded from NCBI GenBank and imported to the KBase platform by remote undergraduate researchers using their respective SRA numbers (Table S1) (Arkin et al., 2018). Sample metadata associated with each metagenome was obtained from Table 1 of Ward et al (2021). Briefly, these metagenomes represent unialgal polycultures containing primarily Cyanobacteria and their cyanosphere heterotrophic microbial associates collected from three aquatic and 47 terrestrial habitats across four broad biomes (arid: 28, tropical: 16, continental: 3, temperate: 3). The dominant Cyanobacterial members in these cultures are phylogenetically diverse spanning five taxonomic orders (Nostocales: 21, Pseudanabaenales: 12, Oscillatoriales: 7, Chroococcales: 5, Synechococcales: 3, Pleurocapsales: 2). These Cyanobacteria-dominant co-cultures were isolated by various phycologists between 1965-2015 and have been maintained in two culture collections at John Carroll University (JCU) and University of Nevada - Las Vegas (UNLV) on solid or liquid Z8 media (an oligotrophic freshwater algae medium (Carmichael, 1986)). Although dates of isolation span 50 years, close to identical culture conditions (use of Z8 medium, 16/8 hr light/dark photoperiod, low light intensity, grown in climate controlled chambers) have been maintained at JCU since 1992 which have been reproduced at UNLV. Collection years can be found in Supplemental Table S1 and Ward et al. (2021).

### Identification of viral contigs

Initial viral analysis was performed following the KBase “Viral Analysis End-to-End” workflow (https://kbase.us/n/75811/85/) by remote undergraduate researchers during the COVID-19 pandemic. Sequence quality for each sample was assessed in KBase using FastQC v0.11.5 (Andrews). Afterwards, low-quality regions and residual Illumina adapters were trimmed using Trimmomatic v0.36 (Bolger, Lohse & Usadel, 2014). This step was followed by a reassessment of the quality of the sequence data again using FastQC. Next, the reads were assembled into contigs with MetaSPAdes v3.13.0 with k-mer sizes 21, 33, 55, 77, 99 and 127 (Nurk et al., 2017). Three samples failed to assemble at this stage using MetaSPAdes due to memory constraints in KBase; for these three samples, MEGAHIT v1.2.9 was utilized instead (Li et al., 2015). Subsequently, all contigs shorter than 5000 base pairs were filtered out using the KBase application “Filter Assembled Contigs By Length - v1.1.2”.

Next, VirSorter v1.0.5 was used to predict the viral genomes in each sample and categorize them into six categories based on their predicted length and viral characteristics (Roux et al., 2015). Viral contigs from VirSorter categories 1-2 (high-confidence lytic viruses) and 4-5 (high-confidence proviruses) were merged into one file using the KBase application “Modify Bins in BinnedContigs - v1.0.2” and then extracted using the “Extract Bins as Assemblies from BinnedContigs - v1.0.2” application. The viral contigs for each sample were then exported from KBase. All predicted viral contigs were combined into a single file and then run through VIBRANT v1.2.1 in “virome” mode (Kieft, Zhou & Anantharaman, 2020). This second viral identification step with VIBRANT was performed to ensure proper trimming of proviruses. CheckV v0.8.1 was then run on the resulting file of predicted viral genomes using the “end_to_end” workflow to calculate viral genome quality, completeness and further predict and remove flanking host regions from integrated proviruses (Nayfach et al., 2021). When performing viral operational taxonomic units (vOTU) analyses, we removed any vOTUs that CheckV could not make quality predictions for (i.e. where quality was “not-determined”).

### Clustering of viral contigs across all samples & relative abundance

Viral genomes were then clustered into 95% identity vOTUs using CD-HIT using the following options: -c 0.95 -aS 0.85 -M 0 -d 0 (Li & Godzik, 2006). In order to assign a putative taxonomy to vOTUs, we first used Prodigal v. 2.6.3 to predict open reading frames in vOTU representative genomes using the -p meta option (Hyatt et al., 2010). We then used VContact2 to cluster vOTUs with the INPHARED August 2022 reference database into viral clusters (VCs) (Bin Jang et al., 2019; Cook et al., 2021) and assigned predicted taxonomy to viruses based on cluster membership as in Santos-Medellin et al (Santos-Medellin et al., 2021). For vOTUs in VCs of interest, ViPTree was further used to place vOTUs into proteomic trees with viral reference genomes using ViPTree database v. 3.6 with virus-host information from RefSeq release 218 (Nishimura et al., 2017).

To quantify the relative abundance of vOTUs across samples, we mapped reads from each metagenome to vOTUs using BBMap with a minid=0.90 (Bushnell, 2022). We used SAMtools to convert sam files to bam files and BEDTools genomecov to get coverage information for each vOTU across each sample from the bam files (Li et al., 2009; Quinlan & Hall, 2010). We then used bamM to parse the bam files to calculate the trimmed pileup coverage (tpmean), the average number of reads aligned to each base after removing the highest 10% and lowest 10% of positions, which we used as a proxy for viral relative abundance (“Ecogenomics/BamM: Metagenomics-focused BAM file manipulation”). We then filtered out vOTUs which displayed < 75% coverage over the length of the viral sequence in R v. 4.3.0 (R Core Team, 2021). Thresholds for analysis of relative abundance of vOTUs were based on previous community guidelines for identity (i.e. ≥ 95% identity) and detection (i.e. ≥ 75% of the viral genome length covered ≥ 1x by reads at ≥ 90% average nucleotide identity) (Roux et al., 2017, 2018). Viral relative abundance was visualized in R using tidyverse, phyloseq, GGally, network, patchwork, ggnewscale, ggforce and ggtext (Butts, C, 2008; McMurdie & Holmes, 2013; Wickham et al., 2019; Schloerke et al., 2021; Pedersen, 2022a,b; Wilke & Wiernik, 2022; Campitelli, 2023). Analysis scripts for the post-KBase portions of this work are available on GitHub (https://github.com/stajichlab/Terrestrial_Cyanophage) and archived in Zenodo (Ettinger, Pietrasiak & Stajich).

### Putative identification of hosts

In order to predict possible microbial hosts for the viral genomes identified, we used NCBI BLAST which has been previously reported to be sufficient for preliminary genus-level host identification (Edwards et al., 2016; Kothari et al., 2021). We downloaded all complete bacterial and archaeal genomes, as of April 7, 2022, from NCBI RefSeq using “ncbi-genome-download” (“kblin/ncbi-genome-download: Scripts to download genomes from the NCBI FTP servers”). In total this amounted to 25,984 bacterial and 416 archaeal genomes. We then used “makeblastdb” from NCBI BLAST 2.12.0+ to convert these complete genomes into a blast database and “blastn” to compare vOTUs to this reference (Camacho et al., 2009). We filtered the results of this analysis in R based on accepted thresholds from the literature as follows: matches had to share ≥ 2000 bp region with ≥ 70% sequence identity to the viral genome and possible host matches needed to have a bit score of ≥ 50 and minimum e value of 0.001 (Edwards et al., 2016; Roux et al., 2016; Kothari et al., 2021; Ettinger et al., 2023). As was done in Kothari et al. (2021), only the top five hits matching these thresholds were considered when predicting a host for each viral genome, with host predictions made at each taxonomic level only if all hits were in agreement. Discrepancies in taxonomy between hits resulted in a “mixed” assignment for that taxonomic level. Based on this method, the lowest level of host prediction was genus level, with some predictions only able to be made at higher taxonomic levels (e.g. class or order). For viruses with a predicted Cyanobacterial host and those that were in VCs with reference genomes reported to infect Cyanobacteria, Cenote-Taker 2 v. 2.1.5 (Tisza et al., 2021) was used to annotate these genomes and Proksee (https://proksee.ca/) was used to visualize the resulting annotations (Stothard, Grant & Van Domselaar, 2019). We then used the comparative gene cluster analysis toolbox (CAGECAT) (van den Belt et al., 2023) to run clinker (Gilchrist & Chooi, 2021) to visualize homologous gene clusters between viral genomes and cyanophage reference genomes downloaded from NCBI GenBank (OM373202, BK059895, NC_015463, NC_029032, NC_016564) (Gao, Gui & Zhang, 2012; Huang et al., 2012; Voorhies et al., 2016; Laloum et al., 2022).

## Results and Discussion

### Successful viral identification from Cyanobacterial co-cultures

In total, 814 viral sequences were identified across the 50 Cyanobacterial co-cultures (Table S1, S2), with each metagenome having on average ∼16 viral sequences (range: 1-54). Of the 814 viral sequences, CheckV predicted that 188 were integrated proviruses (Fig. 1A). CheckV assigned viral quality to 712 of the 814 viral genomes with 465 predicted to be low-quality, 208 to be medium-quality, 32 to be high-quality, and 7 to be complete (Fig. 1B). Based on MIUViG standards (Roux et al., 2018) the catalog represents 39 high-quality draft genomes (i.e., CheckV high-quality and complete genomes) and 775 genome fragments. The mean length of predicted viral sequences was 27,594 bp (range: 5,013–199,914 bp), with 652 viral sequences ≥ 10,000 bp, and with an average of ∼35 predicted genes per viral sequence. We clustered these 814 viral sequences at 95% identity into 726 vOTUs, of which 585 vOTUs represented sequences ≥ 10,000 bp in length. These numbers are similar to previous metagenomic viral discovery efforts in cyanolichen metagenomes (Ponsero et al., 2021). While this work provides a catalog of the dominant DNA viruses that infect terrestrial Cyanobacteria and associated cyanosphere heterotrophs, we explored relatively low diversity microbiomes which were maintained in lab culture conditions. It is known that detection of viruses from such low diversity metagenomes only captures a small amount of DNA virus diversity compared to viromes (Santos-Medellin et al., 2021), and we may have missed RNA viruses entirely which are unlikely to appear in DNA-based metagenomes. Further, VirSorter1, which we use for viral identification, was designed to detect DNA phage in the former order Caudovirales (now Caudoviricetes class), and thus this catalog may be biased towards detection of these phages as well (Roux et al., 2015). Finally, given that this catalog represents viruses identified from co-cultures of Cyanobacterial isolates with their associated cyanosphere microbiome, and not environmental samples directly, we are likely sampling a small subset of the microbiomes of the polyextreme terrestrial environments they originated from. Additionally, any reduction in viruses during passaging may lead to further underestimation compared to the original viral diversity occurring in these environments.

**Figure 1:**
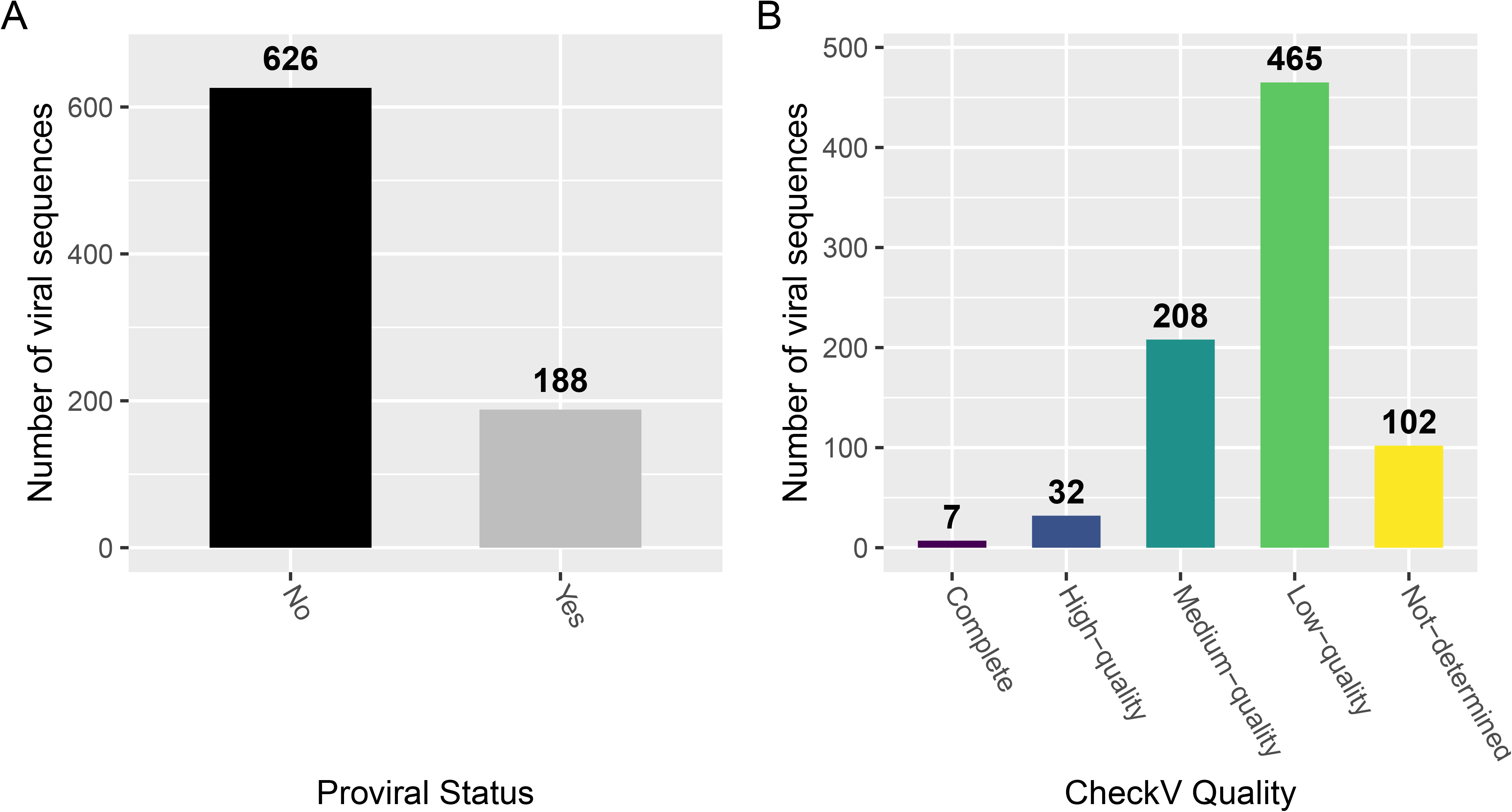
Viral sequences identified from Cyanobacterial co-cultures. (A) The proviral status of predicted viral genomes is shown as a bar graph, with the number above the bar representing the total number of sequences. (B) CheckV quality predictions for viral sequences are shown as a bar graph with bars colored by quality and the number above the bar representing the total number of sequences in each bar.

### Clustering of viral sequences with INPHARED reference data reveals possibly novel genera

To be able to assign putative taxonomy to vOTUs, we further used gene-sharing networks to cluster them with publicly available reference data to form VCs which approximate genus-level groupings. In total 305 vOTUs, representing 341 viral sequences, were placed in 111 VCs. Of these 111 VCs, 26 included representatives from the INPHARED reference dataset, while 79 represented putatively novel genera. Of the 26 VCs that included members that clustered with the reference dataset, 9 VCs included members that were identified as possible proviruses. One VC (VC 1442_0) included a reference viral genome for CrLLKS3, a novel lambda phage in the former Siphoviridae (now unclassified), which was previously reported to infect the aquatic Cyanobacterial genus Cylindrospermopsis (Laloum et al., 2022). The remaining VCs included references viruses reported to infect members of the phyla Proteobacteria (Pseudomonadota; 17 VCs), Spirochaetota (1 VC), Bacteroidota (1 VC), Bacillota (1 VC), Actinomycetota (1 VC), Cyanobacteria (1 VC), or were unreported (4 VCs). The majority of the 79 unique VCs originated from Cyanobacterial co-cultures isolated from arid environments, which is not unexpected as the majority of co-cultures were isolated from arid environments (56% of co-cultures) (Fig. 2A-B). This limited overlap with existing reference data is similar to recent work on cyanolichen metagenomes, where only 28 of 133 VCs included reference genomes (Ponsero et al., 2021). This is different from some metagenomic reports of phage from Cyanobacteria-dominated aquatic systems, including hot springs and lakes, where the majority of reported phage sequences were related to reference data (Skvortsov et al., 2016; Guajardo-Leiva et al., 2018a; Potapov et al., 2019). However, other efforts in freshwater and marine environments have reported a more similar percentage of unclassified viruses compared to reference databases (Sieradzki et al., 2019; Kavagutti et al., 2019). As viral exploration of Cyanobacteria-dominated and other extreme environments continues, we expect to see more sequences clustering with reference genomes over time.

**Figure 2:**
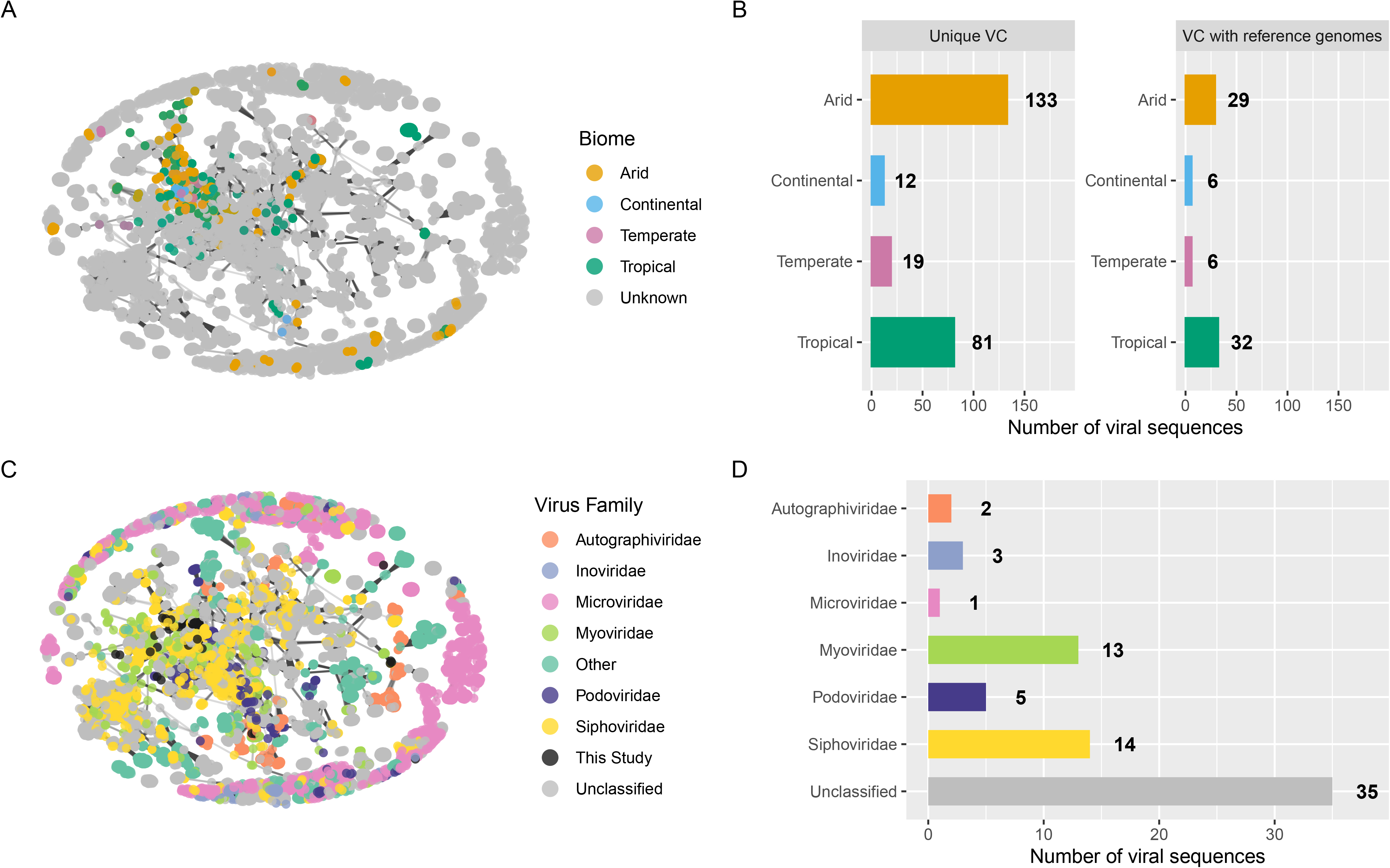
Viral catalog comparison to reference databases (A) Gene-sharing network of vOTUs that formed viral clusters (VCs), including VCs unique to this dataset. Each node is a viral sequence and is colored by the broad biome of the Cyanobacterial co-culture (sample sizes: arid: 28, tropical: 16, continental: 3, temperate: 3). Edges connect sequences with overlap in predicted protein content. (B) Bar charts displaying the number of viral sequences in VCs from this study colored by biome and shown facetted by whether the VC clustered with reference data. Bars are colored by biome, as in A, and the number next to the bar represents the total number of sequences it represents. (C) Subnetwork of VCs with members that include reference genomes obtained from INPHARED. Each node is a viral sequence and is colored by the predicted historical viral taxonomic family for the VC, with the exception of sequences identified in this study which are colored in black. Edges connect sequences with overlap in predicted protein content. (D) Bar chart displaying the number of viral sequences from this study assigned to historical viral families based on VC membership with reference genomes from INPHARED. Bars are colored by historical viral families, as in C, and the number above the bar represents the total number of sequences it represents. For A-D, only viral sequences generated in this study where CheckV was able to predict quality are shown.

The 26 VCs that clustered with the reference data represented 75 of the 814 predicted viral sequences in this dataset. Following new taxonomic recommendations, many viral reference genomes have not yet been reclassified resulting in the majority of the 26 VCs being unclassified under the new schema (Turner, Kropinski & Adriaenssens, 2021). Under the old schema, the majority of VCs (15) were clustered with members assigned to the Caudovirales order (now Caudoviricetes class) of tailed double-stranded DNA viruses, which have also been observed to be the dominant viruses in cyanolichens, biocrusts, hypoliths, and aquatic Cyanobacteria-dominated metagenomes (Sabehi et al., 2012; Zablocki et al., 2014; Adriaenssens et al., 2015; Zablocki, Adriaenssens & Cowan, 2016; Skvortsov et al., 2016; Zeigler Allen et al., 2017; Guajardo-Leiva et al., 2018b; Van Goethem et al., 2019; Ponsero et al., 2021). Within this, VCs were further classified as belonging to the still accepted families Autographiviridae, Inoviridae, and Microviridae, as well as to the recently abolished families Myoviridae, Podoviridae, and Siphoviridae (now unclassified) (Fig. 2C-D). The former Myoviridae, Podoviridae and Siphoviridae represent the main viral families that have historically included cyanophages isolated from marine systems (Mann et al., 2005; Sullivan et al., 2005, 2009, 2010; Weigele et al., 2007) However, we cannot assume all members of VCs classified as belonging to these historical families here are cyanophage as these families also are known to contain members that do not infect Cyanobacteria.

Only 15 of the 39 sequences representing high-quality or complete viral genomes, which were made up of 14 vOTUs, came together to form 13 VCs. Of these, only five VCs included representative viruses from the INPHARED reference data, while 8 VCs were potentially novel genera. Of the VCs that clustered with reference genomes, one was clustered with the order Petitvirales, one with the order Tubulavirales, and three with members of the former order Caudovirales (now Caudoviricetes class). Additionally, VCs included references viruses reported to infect members of the phyla Proteobacteria (Pseudomonadota; 4 VCs), or were unreported (1 VC).

### Most abundant vOTUs are unclassified or unclustered

While the number of viral sequences recovered from each Cyanobacterial co-culture varied (Fig. 3A), we were also curious about the distribution of viral sequences across co-cultures. Thus, we mapped metagenomic reads from each sample to the entire catalog of vOTUs to calculate their relative abundance across all co-cultures. Of 726 vOTUs, only 617 vOTUs had quality scores from CheckV and were detected above thresholds set for inclusion. Looking at the relative abundance of these remaining 617 vOTUs, we found that the majority of co-cultures were dominated by unclustered vOTUs or by vOTUs that formed VCs that did not cluster with INPHARED reference genomes (Fig. 3B). Only a small portion, on average 8.2% (range: 0 – 100%), of vOTU relative abundance in samples was from vOTUs in VCs that included reference genomes. The majority of vOTUs, 496 (80.4%), were singletons and only found in one sample. While a total of 121 vOTUs and 86 VCs (13 of which clustered with reference data) were detected in two or more samples. The most prevalent VC (714_0) did not cluster with INPHARED reference genomes and represented 25 vOTUS across 26 samples spanning different habitats and collection locations. Using ViPTree, we found that vOTUs in this VC form a unique monophyletic clade within the recently abolished family, Siphoviridae (now unclassified) (Fig. S1). Overall these results demonstrate how few viruses are present in reference databases relative to existing viral diversity in Cyanobacteria-dominant co-cultures and highlight the importance of viral discovery efforts in addressing this knowledge gap. Low diversity co-cultures from public or research culture collections, such as those investigated in our study, could function as manageable study systems to establish viral reference benchmarks in future comparative works.

**Figure 3:**
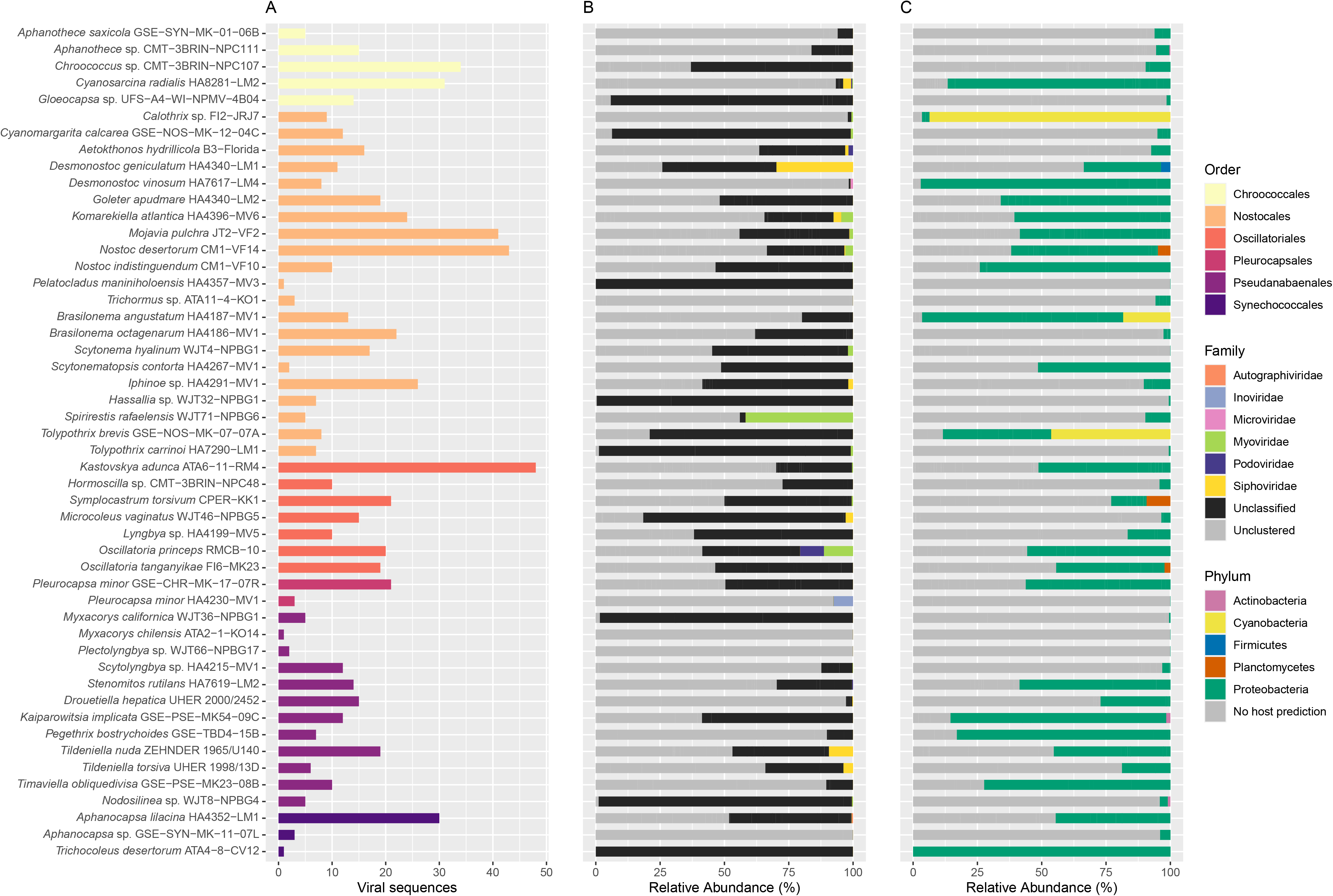
Viral communities across Cyanobacterial co-cultures. (A) The number of viral sequences predicted from each co-culture metagenome. Bars are arranged and colored based on the taxonomic order the Cyanobacteria in the co-culture is taxonomically classified as. (B) Stacked bar charts displaying the taxonomic relative abundance of vOTUs in each co-culture. Bars are colored based on predicted historical viral taxonomic families. Viral sequences that clustered into VCs, but that lack taxonomy (i.e., unique VCs that lack reference data, or VCs where reference data taxonomy is unclear) are labeled “Unassigned”, while viral sequences that did not cluster into VCs are labeled “Unclustered”. (C) Stacked bar charts of the relative abundance of vOTUs based on predicted hosts in each co-culture. Bars are colored based on the predicted host taxonomy, at the phylum level, of each vOTU . For A-C, only viral sequences where CheckV was able to predict quality are shown.

### Proteobacteria (Pseudomonadota), not Cyanobacteria, dominate host predictions

We explored possible host associations by using NCBI BLAST to compare vOTUs against a collection of complete bacteria and archaea genomes from RefSeq. This method has been reported to have high accuracy, but low recall as it is limited by existing databases (Coclet & Roux, 2021). The majority of viral sequences (65.8%) had no host prediction. However, we were able to predict possible bacterial hosts for 211 out of 617 vOTUs to at least the phylum-level (Fig. 3C). Of these 211 predictions, we were able to classify the hosts for 151 vOTUs to genus-level with the most common predicted host genera being *Brevundimonas* (15 vOTUs), *Sphingomonas* (13), *Pseudomonas* (11), *Bradyrhizobium* (11) and *Bosea* (10). Many of these genera have been reported in association with Cyanobacteria from both aquatic and terrestrial habitats (Bruno et al., 2006; Shi et al., 2009; Li et al., 2012; Kim et al., 2019). For those viruses with a host prediction, zooming out these were largely projected to infect Proteobacterial (now Pseudomonadotal) hosts (95.3%). Within the Proteobacteria (Pseudomonadota), almost half (47.3%) of the predicted hosts were members of the order Hyphomicrobiales. Further, 18 of the 25 vOTUs represented in the most shared VC (714_0) were predicted to infect members of the order Hyphomicrobiales.

Only three vOTUs (vOTU602, vOTU638, vOTU669), representing three unclassified viral sequences, had Cyanobacterial host predictions, and none of these belonged to the single VC (VC 1442_0) that included reference virus members reported to infect Cyanobacteria. Of these three vOTUS, two were predicted to be medium-quality (vOTU602, vOTU669) and one complete (vOTU638). Two of the three vOTUs formed separate VCs unique to this data (vOTU602 = VC 1342_0; vOTU638 = VC 1447_0). All three vOTUs were predicted to have hosts in the order Nostocales. However for two vOTUs we were able to only predict hosts to the order-level (vOTU602, vOTU638), while the third vOTU was predicted to infect the genus *Calothrix* (vOTU669). These predictions match the taxonomy of the Cyanobacterial member of the cultures that they were obtained from. Two of these vOTUs (vOTU602, vOTU669) originated from cultures isolated from desert environments, while the third originated from a culture isolated from tropical soil (vOTU638). Each of these three vOTUs was only detected in the culture it originated from. Using ViPTree, we found that these vOTUs were placed in a clade where many viral sequences had aquatic Cyanobacterial host predictions (Fig. S2). However the vOTUs were on long branches with limited similarity to reference genomes and no reference viruses were close relatives to the three vOTUs here. While finding only three cyanophage vOTUs was surprising given the predominance of Cyanobacteria in these cultures, previous work from biocrusts and hypoliths have similarly reported a low abundance of cyanophage (Zablocki, Adriaenssens & Cowan, 2016; Van Goethem et al., 2019). An important caveat to remember is the reliance of host predictions on available genomes; as Cyanobacterial genome availability increases, we expect that ability to link viruses to their hosts in these ecosystems will increase as well.

Despite not having a Cyanobacterial host prediction using BLAST, we also used ViPTree to investigate the two vOTUs in VC 1442_0 (the VC that included the reference virus, CrLLKS3, reported to infect aquatic Cyanobacteria). The vOTUs in VC 1442_0 represent low-quality (vOTU185) and medium-quality (vOTU15) viral genomes from cultures isolated from aquatic plant and cave rock wall environments respectively. Using ViPTree, we found that vOTU15 was placed on a long branch in a clade that included CrLLKS3 and it’s closest relative Cy-LDV1 (Laloum et al., 2022), but no other viruses that had Cyanobacterial host predictions (Fig. S3). However, CrLLKS3 was reported to have higher genomic similarity to heterotrophic bacteria phage than known cyanophage (Laloum et al., 2022), so it is possible these vOTU15 may just represent another member of this novel cyanophage group. Despite being in the same VC, vOTU185 was placed elsewhere in the tree. We believe this is an erroneous placement due to vOTU185’s low-quality and small size.

In an attempt to gain further insight about the possible interactions of these five vOTUs with their possible Cyanobacteria hosts, we annotated these viral sequences using Cenote-Taker2 to look for putative candidate AMGs. Cyanophages in aquatic ecosystems often are reported to have AMGs related to photosynthesis, phosphorus utilization, and carbon metabolism (Bailey et al., 2004; Sullivan et al., 2005, 2010; Millard et al., 2009). However, in all five vOTUs, we found the majority of non-viral genes to be unannotated (i.e., hypothetical proteins) and observed no clear AMG candidates related to common cyanophage functions (Fig.S4). Further assessment of homologous gene clusters between vOTUs and cyanophage close relatives, based on ViPTree analyses, did not identify any homologs that shared > 50% protein similarity. Instead, we mainly observed lower similarity matches to viral hallmark genes or hypothetical proteins (Fig. S5). One reason for the lack of clear AMG candidates may relate to viral completeness. Four of the five vOTUs were predicted by CheckV to be low or medium-quality, thus the inability to detect AMGs may relate to these viral sequences being incomplete. Another possibility is that the lack of AMG candidates may be representative of phage lifestyle (lytic vs. temperate), with lytic phage more likely to focus on hijacking host replication machinery and less driven to promote long-term host fitness through AMGs (Breitbart et al., 2018; Warwick-Dugdale et al., 2019; Luo et al., 2022). Regardless, as more studies focus on viral discovery and characterization efforts, the power to identify and annotate AMGs will increase leading to better overall comprehension of the role of viruses in Cyanobacteria-dominated ecosystems.

## Conclusion

To expand knowledge of viruses infecting Cyanobacteria-dominated microbiomes, we employed viral discovery methods on existing metagenomes from nonaxenic Cyanobacterial co-cultures as part of remote undergraduate research experiences (Ward et al., 2021). We were able to identify 814 viral sequences including sequences that formed unique VC and which may represent novel viral genera. While this work provides a catalog of viruses that infect terrestrial Cyanobacteria and associated cyanosphere heterotrophs, due to methodological limitations, we are likely only sampling DNA phage and are undersampling viruses in these cultures. Future work should seek to profile terrestrial Cyanobacteria viruses using more comprehensive methods such as through viromics sequencing of environmental samples. While we observed no wide scale biome or Cyanobacterial host patterns driving viral diversity, this is likely due to the limited nature of this work including the incomplete sampling of the viral community, the mixed nature of the nonaxenic cultures, and the fact that these are isolated cultures, not true environmental samples. However despite study limitations, profiling the viral community of co-cultures resulted in our ability to identify viral diversity on par with environmental metagenomic studies, as well as enabled the identification of possible novel viruses, including several potential cyanophage. Overall, future studies should build on this work to include a more extensive investigation of viral-Cyanobacteria interactions in extreme environments and profile the auxiliary metabolic genes of viral genomes which may play roles in various cycling processes in terrestrial Cyanobacteria-dominated environments.

## Acknowledgements

We thank Jeffrey R. Johansen for being a lifelong mentor, collaborator, and friend to N.P. Jeff initiated a culture collection of terrestrial cyanobacteria at JCU from which subcultures of at least half of the investigated strains were obtained and investigated in Ward et al. (2021).

