## Supplemental Table Legends and Supplemental Figures 1-5 for "Cataloging Viral Diversity from Nonaxenic Terrestrial Cyanobacteria Cultures"

**Supplemental Table Legends and Figures:**

**Table S1. Sample information and viral identification statistics from Cyanobacterial co-culture metagenomes.** Here we provide information for each Cyanobacterial co-culture metagenome obtained from Table 1 of Ward et al (2021) including Cyanobacterial species, reported habitat and SRA Accession numbers. We have expanded on the reported data with Köppen climate classifications (which we used for visualization in this study), habitat substrates, as well as the taxonomic order of the expected Cyanobacterial member. This table also reports for each metagenome the number of predicted viral sequences, the number of proviral sequences, the average viral sequence length, average number of genes per sequence and the number of viral sequences identified by CheckV as being Complete, High-quality, Medium-quality or Low-quality.

**Table S2. Viral catalog sequence information.** Here we provide information for each viral sequence in the total catalog of 814 putative sequences including the associated Cyanobacterial species metagenome, viral sequence length, proviral status, gene count, CheckV quality score, vOTU assignment, VC assignment, putative historical viral taxonomy, putative viral taxonomy under new ICTV guidelines, and taxonomy of the predicted host. The three predicted cyanophage vOTUs are highlighted in bold.

**Figure S1: ViPTree depicts VC 714_0 as a monophyletic clade.** The 25 vOTUs in VC 714_0 were placed into proteomic trees with viral reference genomes from RefSeq release 218 using ViPTree. Here we show a subset of the full tree from ViPTree based on highest similarity scores to the vOTUs of interest, with the 25 vOTUs in VC 714_0 shown in the tree in red and further indicated with a red star.

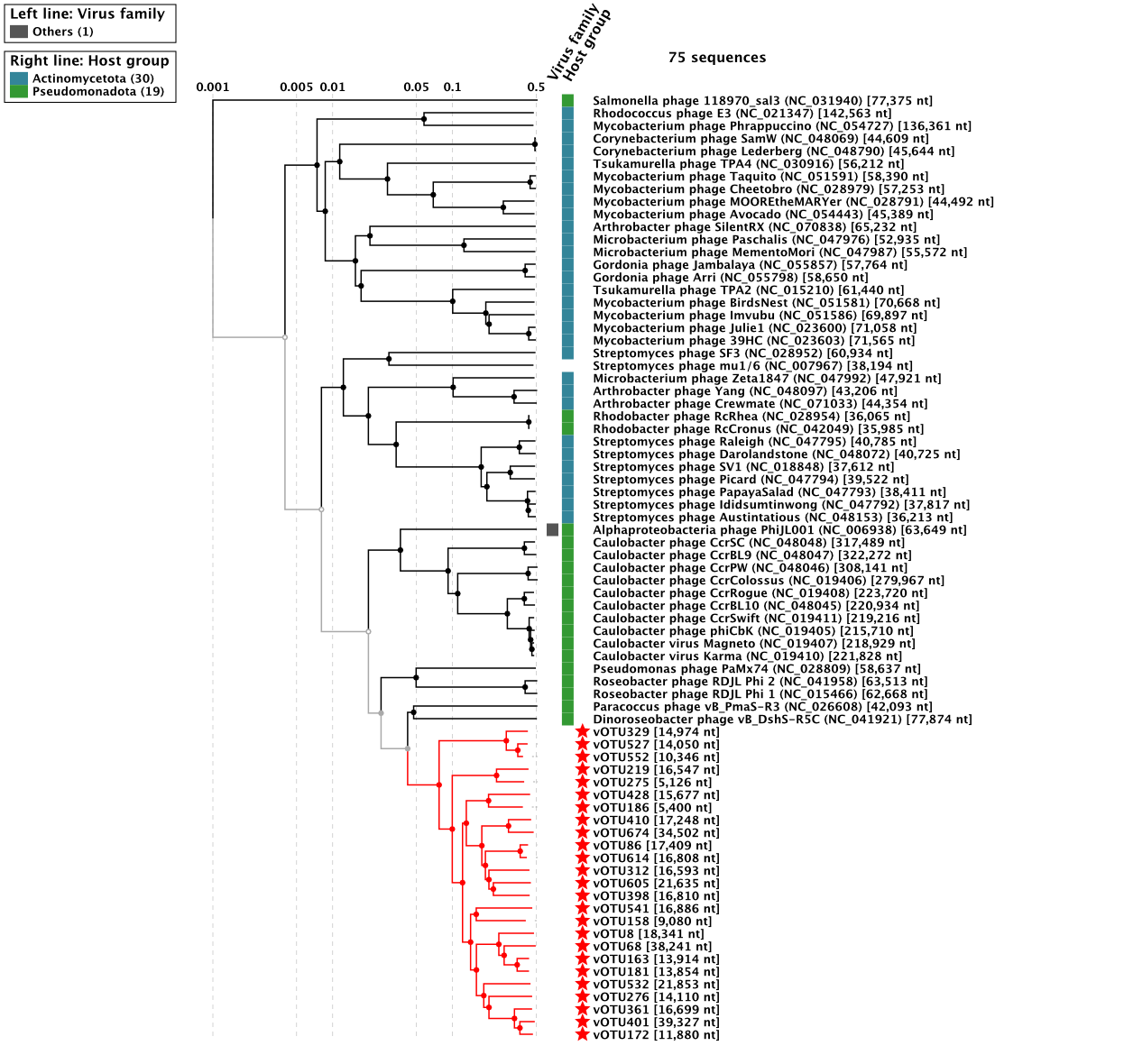

**Figure S2: ViPTree depicts predicted cyanophage in a clade that includes known cyanophages.** The 3 vOTUs (vOTU602, vOTU638, vOTU669) predicted to infect Cyanobacteria were placed into proteomic trees with viral reference genomes from RefSeq release 218 using ViPTree. Here we show a subset of the full tree from ViPTree based on highest similarity scores to the vOTUs of interest, with the 3 vOTUs shown in the tree in red and further indicated with a red star. For all three of these vOTUs, there were no reference genomes with sequence similarities > 0.02 which is ViPTree’s default threshold for “related” genomes.

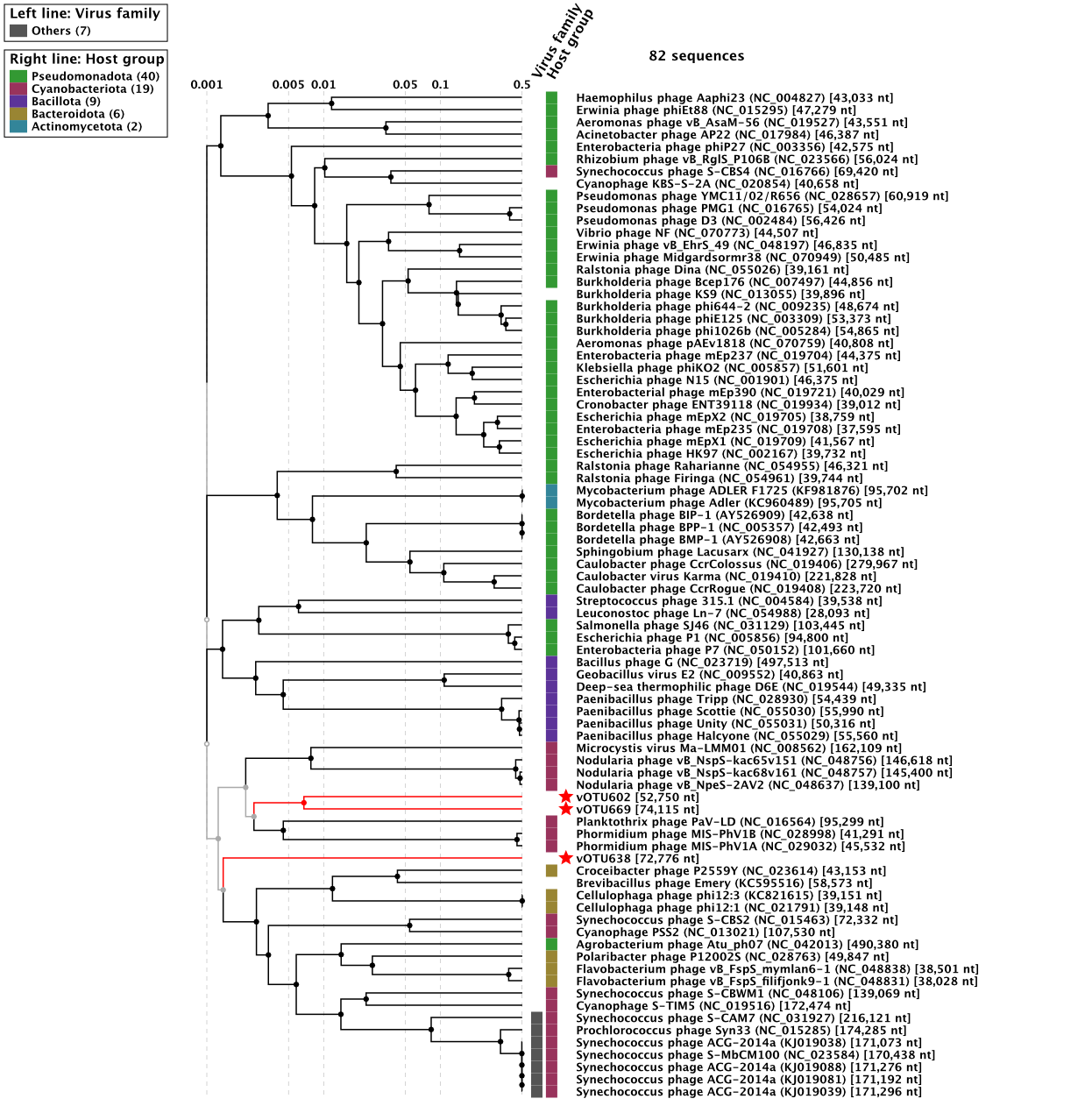

**Figure S3: ViPTree of VC 1442_0.** The three viral genomes in VC 1442_0, which included the reference genome, Cr-LKS3 which infects an aquatic Cyanobacteria, and 2 vOTUs (vOTU15, vOTU185), were placed into proteomic trees with viral reference genomes from RefSeq release 218 using ViPTree. A close relative of Cr-LKS3, Cy-LDV1, was also included. Here we show a subset of the full tree from ViPTree based on highest similarity scores to the vOTUs of interest, with the 2 vOTUs shown in the tree in red and further indicated with a red star. Cr-LKS3 and Cy-LDV1 are show in the tree in orange and further indicated with an orange star.

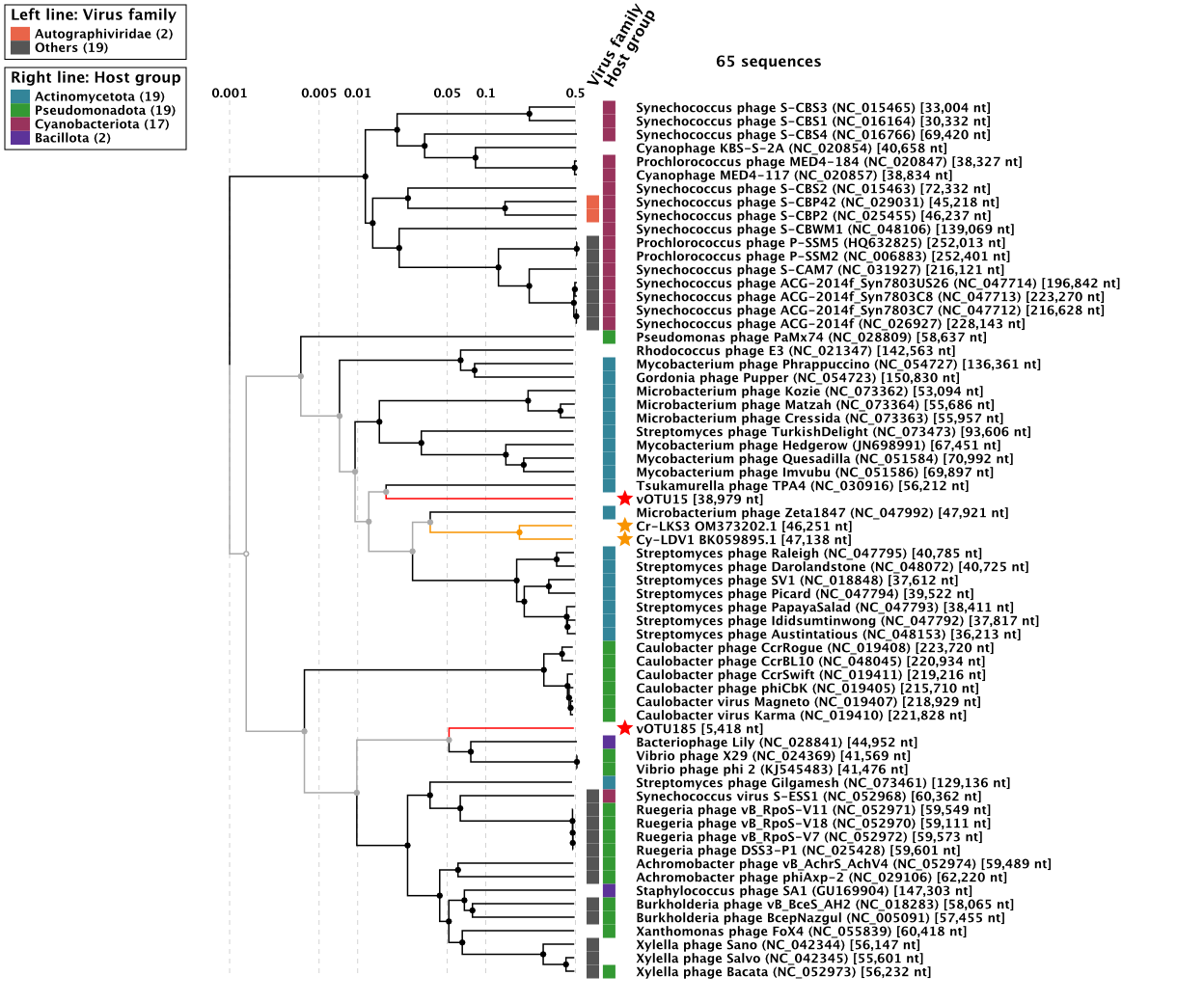

**Figure S4: Genome maps of predicted cyanophages.** The 3 vOTUs (vOTU602, vOTU638, vOTU669) predicted to infect Cyanobacteria and the 2 vOTUs (vOTU15, vOTU185) in VC 1442_0, which included the reference genome reported to infect Cyanobacteria, were annotated using Cenote-Taker2 and visualized using Proksee. Here we depict the genomic architecture of these viral sequences, with GC content on the inside and predicted genes separated by strand on the outside, for (A) vOTU602, (B) vOTU638, (C) vOTU669, (D) vOTU15, and (E) vOTU185. For all 5 vOTUs, viral hallmark genes (as determined by Cenote-Taker 2) are shown in dark purple, viral related genes (based on manual assessment) are shown in light purple, hypothetical proteins are shown in gray, and all other annotated genes are shown in green. While all five vOTUs are shown in circular orientation for visualization purposes, only vOTU 669 was predicted to be circular by Cenote-Taker2.

**(A)**

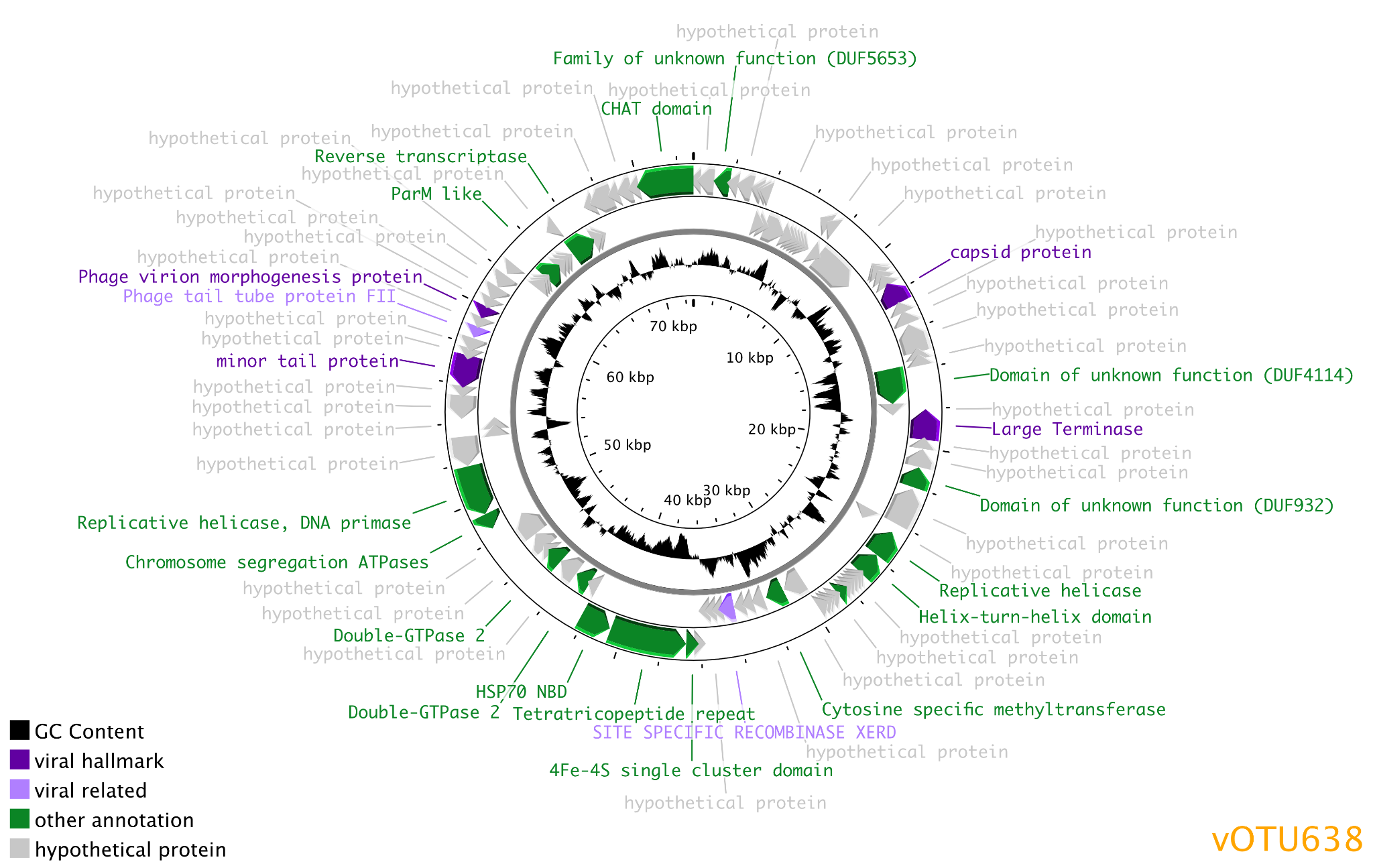

**(B)**

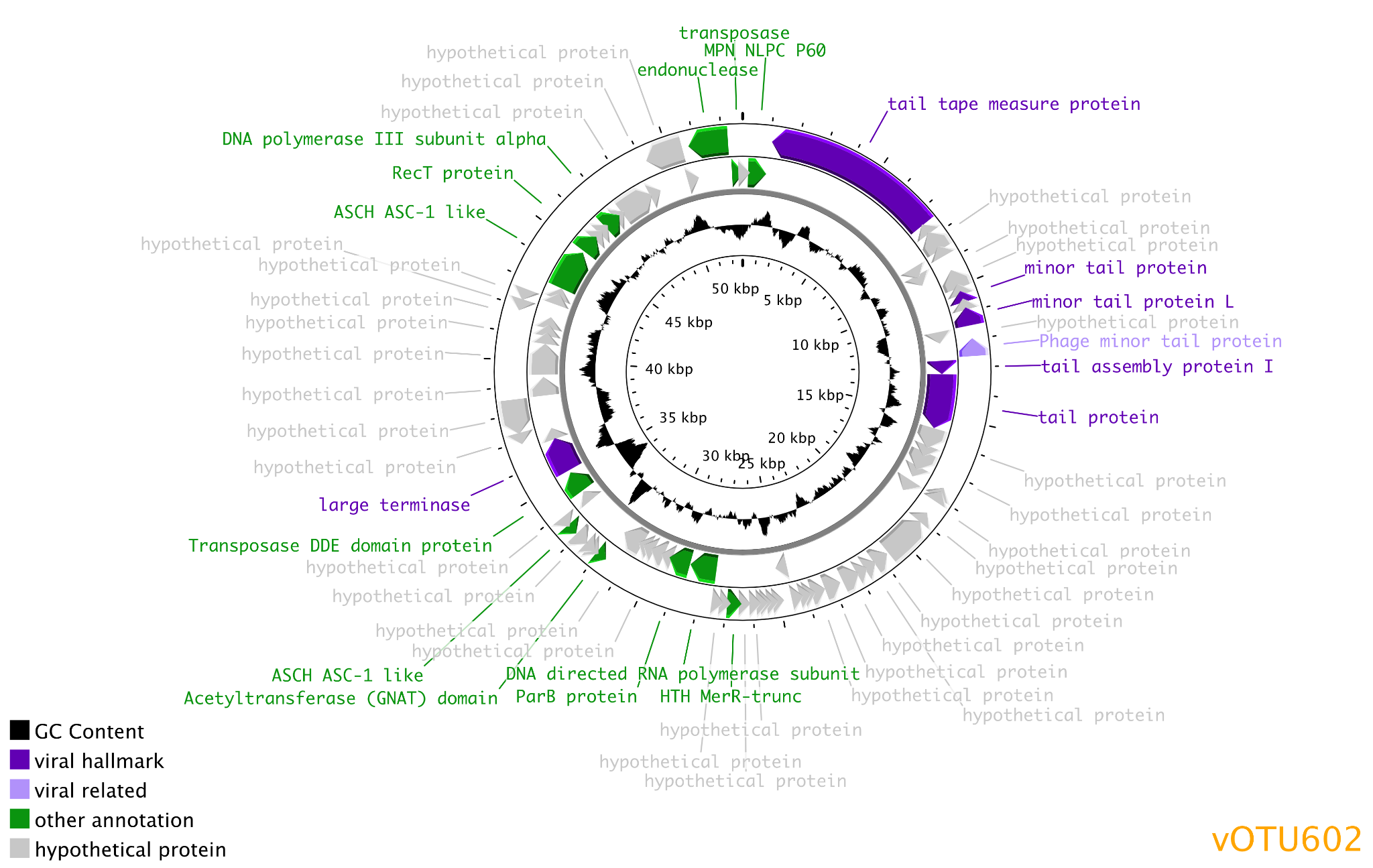

**(C)**

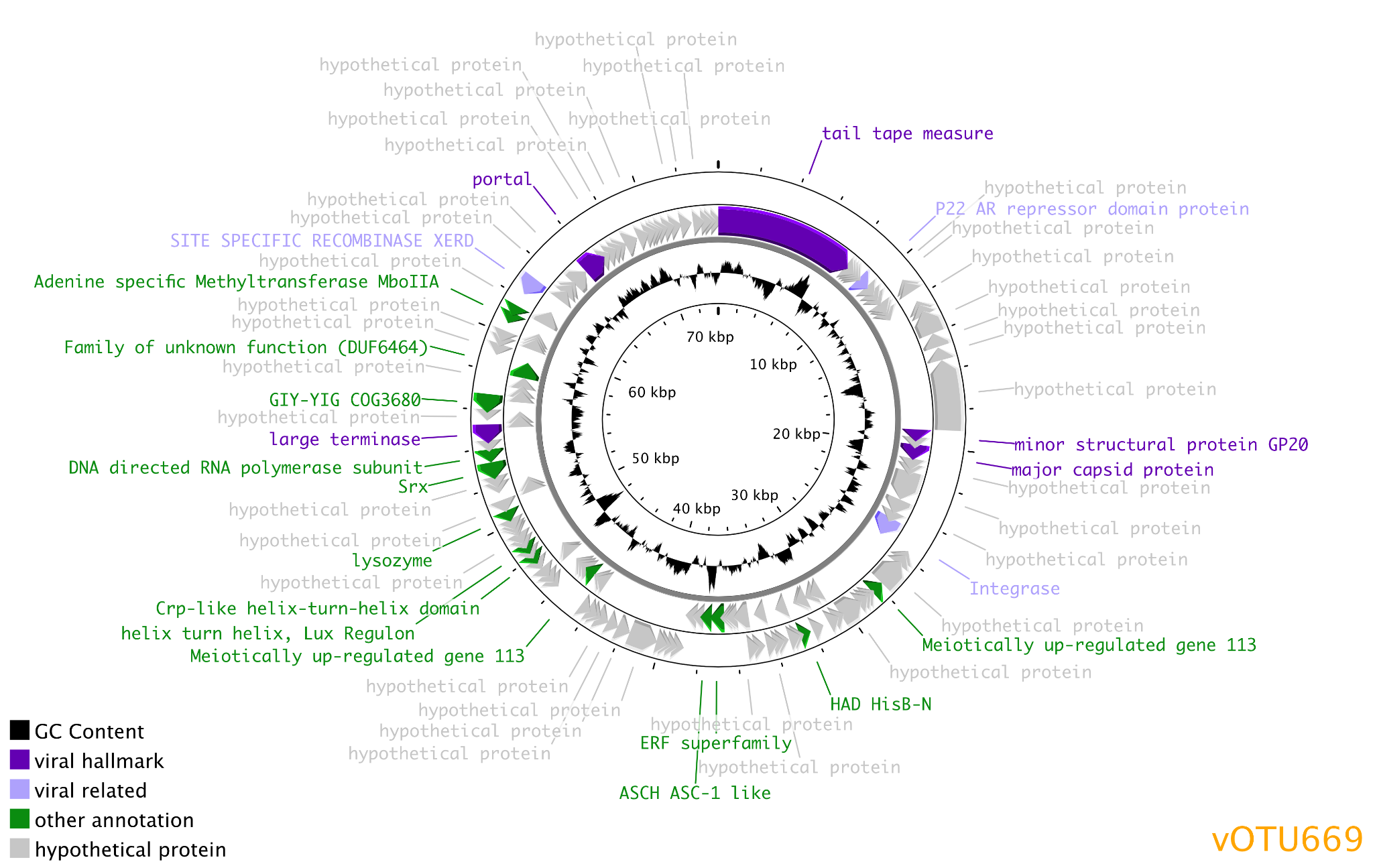

**(D)**

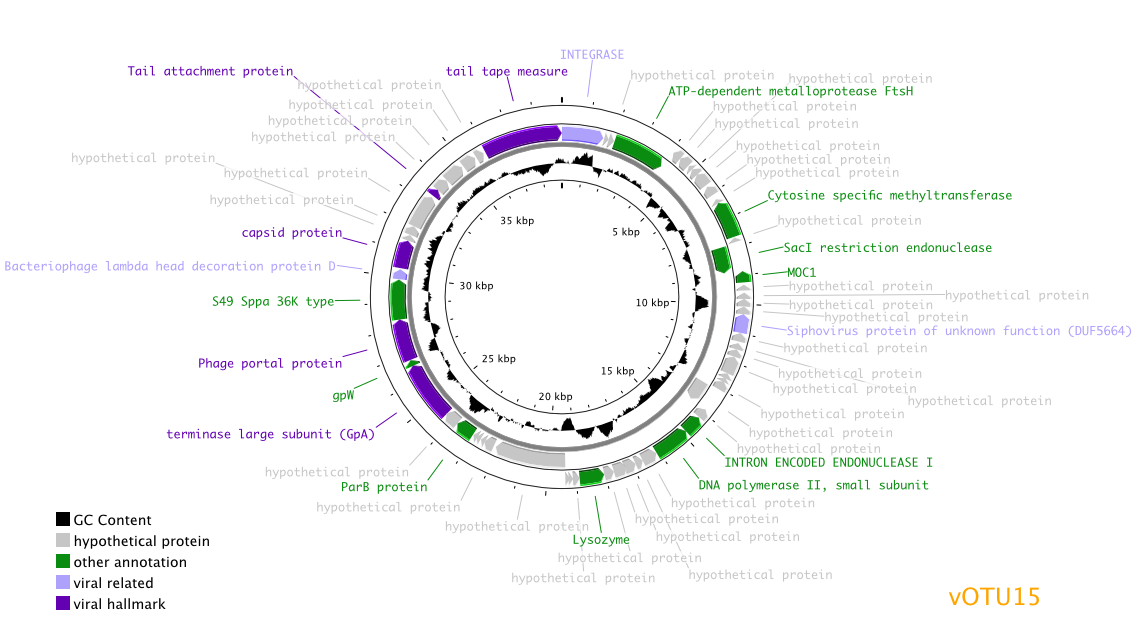

**(E)**

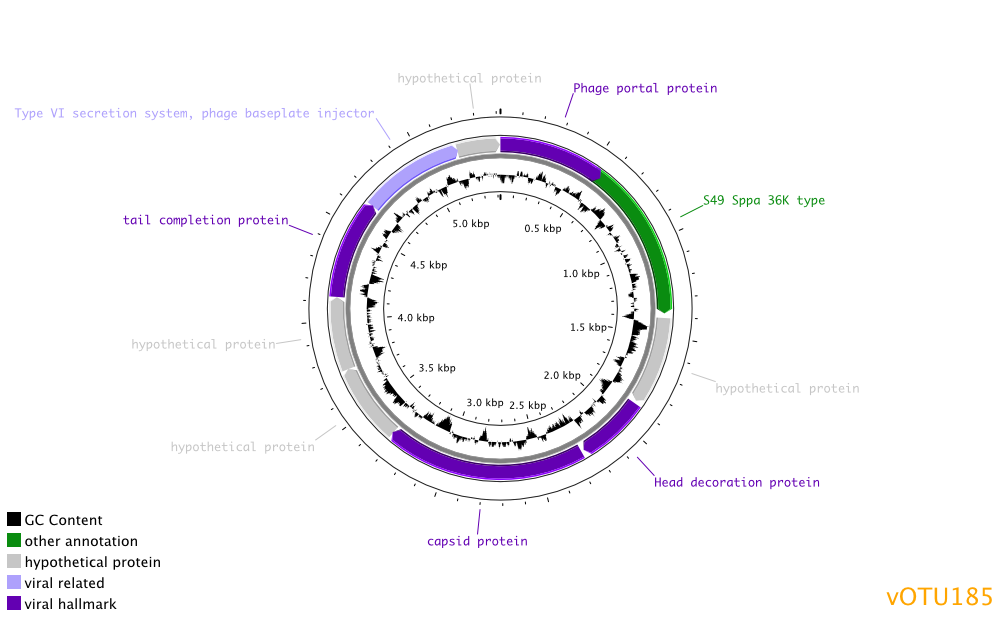

**Figure S5: Homologous gene clusters between vOTUs and known cyanophages.** The 3 vOTUs (vOTU602, vOTU638, vOTU669) predicted to infect Cyanobacteria and the 2 vOTUs (vOTU15, vOTU185) in VC 1442_0, which included the reference genome reported to infect Cyanobacteria, were annotated using Cenote-Taker2. Genomes and annotations of cyanophage close relatives (as determined by VipTree) were then downloaded from NCBI GenBank. CAGECAT was used to visualize homologous gene clusters between vOTUs and reference cyanophage genomes. Here we show homologous gene clusters identified with ≥ 25% protein similarity for (A) vOTU602 and vOTU669, (B) vOTU638, and (C) vOTU15 and vOTU185. Linkages between genes are colored by percent similarity.

**(A)**
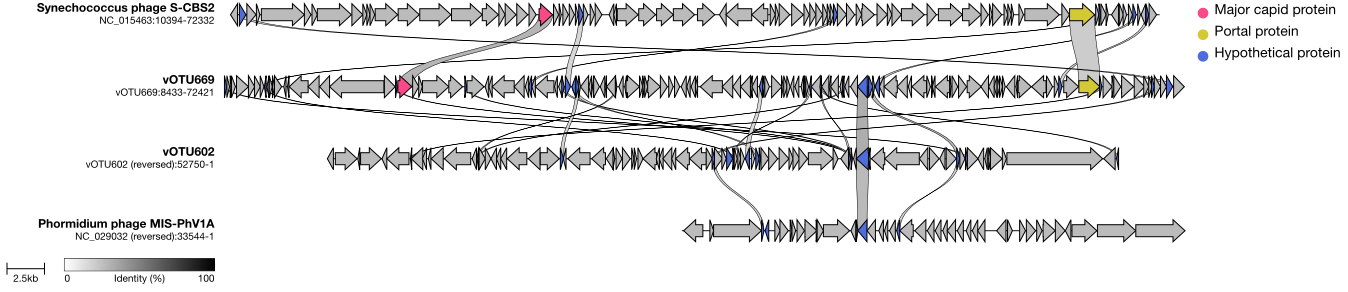

**(B)**

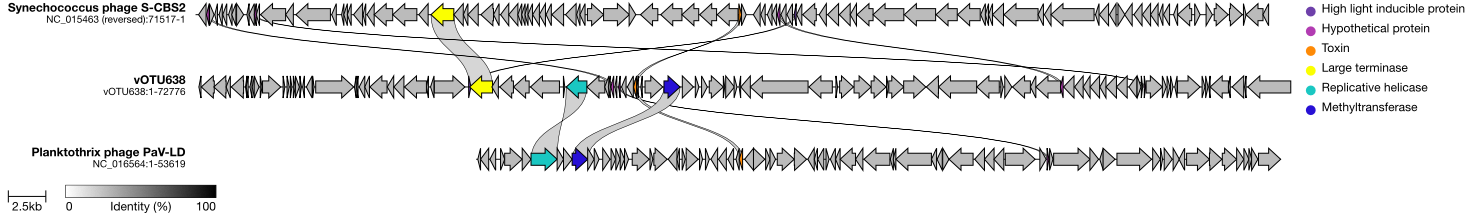

**(C)**

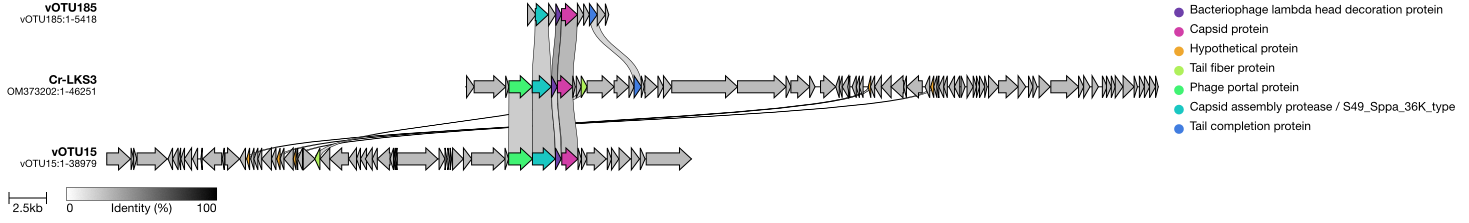
